# Using a modular massively parallel reporter assay to discover context-dependent regulatory activity in type 2 diabetes-linked noncoding regions

**DOI:** 10.1101/2023.10.08.561391

**Authors:** Adelaide Tovar, Yasuhiro Kyono, Kirsten Nishino, Maya Bose, Arushi Varshney, Stephen C.J. Parker, Jacob O. Kitzman

**Affiliations:** Gilbert S. Omenn Department of Computational Medicine and Bioinformatics, University of Michigan, Ann Arbor, MI 48109; Department of Human Genetics, University of Michigan, Ann Arbor, MI 48109; Department of Biostatistics, University of Michigan, Ann Arbor, MI 48109; Current affiliation: Seattle Hub for Synthetic Biology, Allen Institute, Seattle, WA 98109

**Keywords:** transcription factors, enhancers, diabetes, gene regulation

## Abstract

Most genome-wide association signals for complex disease reside in the noncoding genome, where defining function is nontrivial. MPRAs (massively parallel reporter assays) offer a scalable means to identify functional regulatory elements, but are typically conducted without regard to cell type, pairing cloned fragments with a generic housekeeping promoter. To explore the context-sensitivity of MPRAs, we screened enhancer activity across a panel of nearly 12,000 198-bp fragments spanning over 300 type 2 diabetes- and metabolic trait-associated regions in the 832/13 rat insulinoma beta cell line, a relevant model of pancreatic beta cells. We explored these fragments’ context sensitivity by comparing their activities when placed up- or downstream of a reporter gene, and in combination with either a synthetic housekeeping promoter (SCP1) or a more biologically relevant promoter corresponding to the human insulin (*INS*) gene. We identified clear effects of MPRA construct design on enhancer activity. Specifically, a subset of fragments (n = 702/11,656) displayed positional bias, evenly distributed across up- and downstream preference. Promoter choice also influenced MPRA activity (n = 698/11,656), mostly biased towards the cell-specific *INS* promoter (73.4%). To identify sequence features associated with promoter preference, we used Lasso regression with 562 genomic annotations and discovered that fragments with *INS* promoter-biased activity are enriched for HNF1 motifs. HNF1 family transcription factors are key regulators of glucose metabolism disrupted in maturity onset diabetes of the young (MODY), suggesting genetic convergence between rare coding variants that cause MODY and common T2D-associated regulatory regions. We designed a follow-up MPRA containing HNF1 motif-enriched fragments and observed several instances where deletion or mutation of HNF1 motifs disrupted the *INS* promoter-biased enhancer activity, specifically in the beta cell model but not in a skeletal muscle cell line, another diabetes-relevant cell type. Together, our study suggests that cell-specific regulatory activity is partially influenced by enhancer-promoter compatibility and indicates that careful attention should be paid when designing MPRA libraries to capture context-specific regulatory processes at disease-associated genetic signals.

## Introduction

Complex diseases arise from individual and interactive effects of genetic, environmental, and lifestyle factors. Recent genome-wide association studies have revealed that the majority of complex disease-associated loci (∼90%) are found in noncoding regions where they are expected to alter gene regulation (Watanabe et al. 2019). Progress to identify causal variants and define their role(s) in disease pathogenesis has been slow due to the lack of a clear variant-to-function framework for noncoding variants. Massively parallel reporter assays (MPRAs) have emerged as a useful method for screening large collections of putative regulatory elements, learning principles of disease-associated regulatory variation, and efficiently prioritizing candidate causal variants at disease loci (Khetan et al. 2021; Tewhey et al. 2016; Arnold et al. 2013; Bergman et al. 2022).

A wide array of MPRA designs exist, primarily differing by the position where fragments of interest are cloned relative to a basal promoter, and by reporter gene (summarized in (Klein et al. 2020)). Enhancers are classically defined as acting at a distance in an orientation-independent manner (Banerji et al. 1981), though how distinct they are from promoters remains unclear. Klein et al. reported modest impacts of fragment position on enhancer activity, which could nevertheless mask the often subtle regulatory effects being measured (Ulirsch et al. 2016; Choi et al. 2020; Tewhey et al. 2016; Lu et al. 2021; Myint et al. 2020; McAfee et al. 2023), clouding downstream interpretation.

In addition to fragment positioning, another key design consideration is the choice of promoter. To date, most MPRAs used to study disease-associated genetic variants rely on a standard design featuring one of several housekeeping promoters (e.g., minimal promoter [minP], super core promoter [SCP1]) (Tewhey et al. 2016; McAfee et al. 2022; Melnikov et al. 2012; Patwardhan et al. 2012). There is conflicting evidence as to the influence of promoter-enhancer compatibility rules: some studies suggest that unique sequence features including transcription factor motifs dictate how strongly an interaction will occur (Zabidi et al. 2015; Martinez-Ara et al. 2022) while others indicate that there is minimal specificity with a simple additive model combining intrinsic enhancer and promoter strength (Hong and Cohen 2022; Bergman et al. 2022; Sahu et al. 2022). There have been limited comparisons of MPRA design parameters including fragment position and enhancer-promoter compatibility, and how these choices interact at disease-associated loci has not yet been systematically examined. We therefore speculated that using a promoter of a physiologically relevant gene may improve our ability to detect disease-associated regulatory perturbations in a tractable and well-matched cell model.

Here, we used a modular MPRA to assess enhancer activity across multiple promoter and position contexts to evaluate how enhancer-promoter compatibility contributes to context-specific gene regulation in type 2 diabetes.

## Results

### Massively parallel reporter design and cloning

To explore regulatory activity across loci associated with type 2 diabetes (T2D) and related metabolic traits, we designed a library of 13,226 targets spanning regions in high linkage disequilibrium (R^2^ > 0.8) with index signals reported by the DIAMANTE (Mahajan et al. 2022, 2018; Spracklen et al. 2020) and MAGIC (Chen et al. 2021) consortia (**Figure 1**; **Table S1**). We added short 16 bp anchor sequences on both sides of each fragment to enable cloning into the MPRA constructs. We cloned this fragment library into four different MPRA constructs: upstream or downstream of a portion of the human insulin (*INS*) promoter or the synthetic super core (SCP1) promoter (Juven-Gershon et al. 2006) (**Figure S1A**). We simultaneously delivered the resulting plasmid libraries to the 832/13 rat insulinoma cell line via electroporation, then collected cells 24 hours later to isolate RNA. We performed this experiment in triplicate and prepared barcode sequencing libraries from the cDNA and plasmid input to estimate enhancer activity of each fragment (**Figure S1C**).

**Figure 1.**
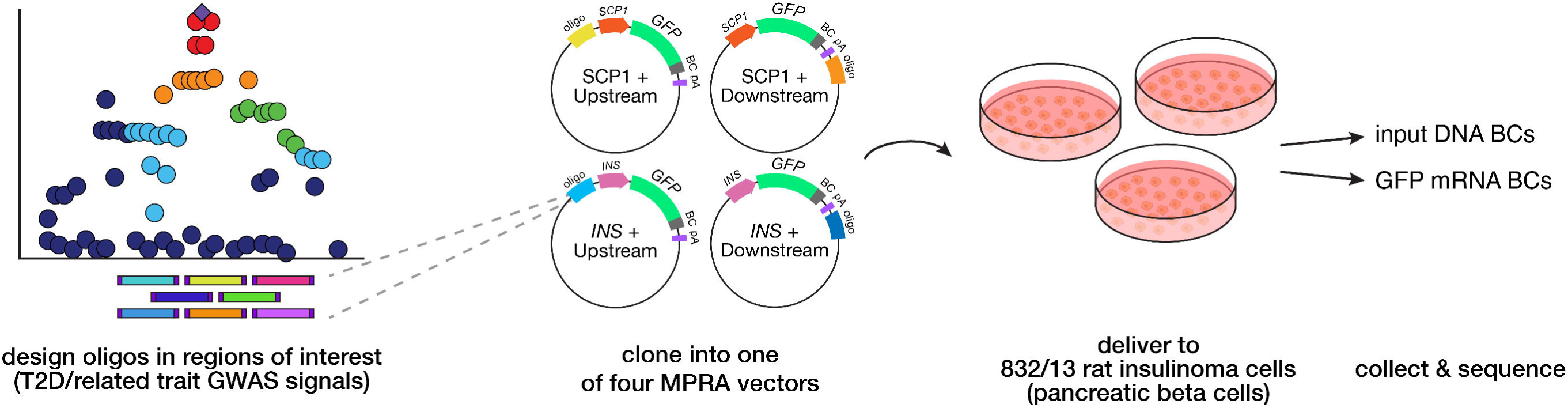
Study design. We synthesized a library of 198-bp fragments encompassing 13,226 sites in high linkage disequilibrium with type 2 diabetes (T2D) and related trait-associated signals. We cloned this library into each of four MPRA constructs: fragment upstream or downstream of the human insulin (*INS*) promoter or upstream or downstream of the super core promoter (SCP1). We simultaneously delivered all four libraries to 832/13 rat insulinoma cells (n = 3) and collected DNA and RNA for sequencing.

### Regulatory activities measured across fragments

Our MPRA library contained both reference and alternate alleles but was not barcoded to sufficient complexity for allelic effect detection. Accordingly, we focused on patterns of regulatory activity across sequence contexts rather than allelic bias in activity. After filtering for fragments represented by greater than two barcodes and with at least ten input DNA counts across all four configurations, we proceeded with a set of 11,656 fragments for downstream analysis.

To qualitatively examine the influence of position and promoter on fragment activity, we used principal component analysis (PCA; **Figure 2A**) on log_2_(RNA/DNA) activity estimates using library size-normalized counts. We identified that the four constructs separate on principal component 2 (13.25% of variance) based on the position into which fragments were cloned, where both downstream constructs cluster together separately from the two upstream constructs. The two upstream constructs further separate on principal component 1 (15.68% of variance) based on the promoter that was used. To evaluate whether other technical factors confounded the group separations we saw using PCA, we calculated pairwise Spearman correlation between the first five principal components, RNA library size, DNA library size, and ratio of RNA library size/DNA library size (library size ratio) (**Figure S2A**). We identified strong negative correlation (R^2^ = 0.972) between PC1 and library size ratio, suggesting a relationship between PC1 and overall enhancer activity for a given library. Additionally, we observed a strong relationship between RNA library size and DNA library size (R^2^ = 0.972), which is expected given that RNA counts are partially dependent on the plasmid DNA counts. Otherwise, we observed no strong correlations between any of the PCs and technical factors.

**Figure 2.**
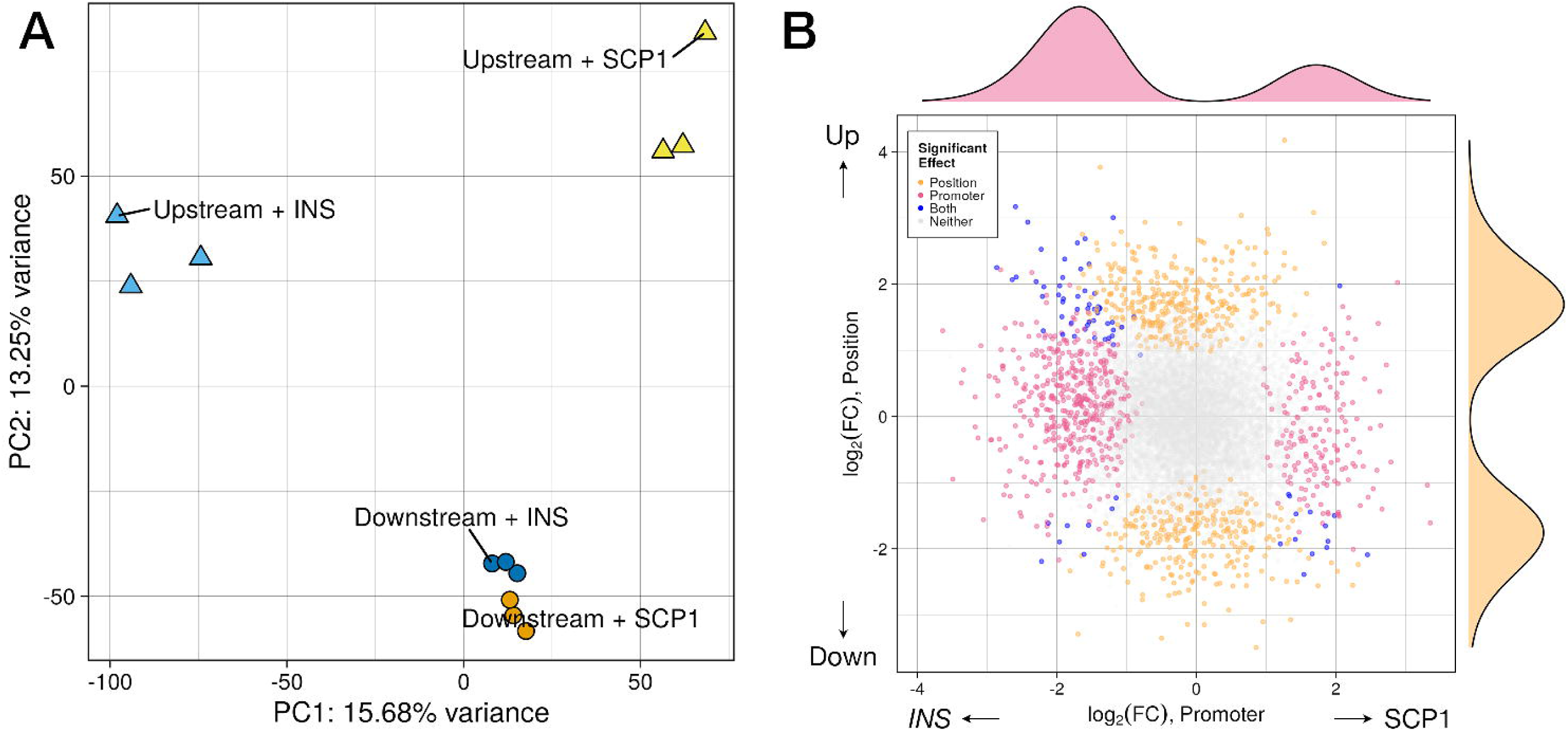
Promoter and fragment position relative to the promoter and reporter gene influence MPRA activity. (A) After fragment activity quantification in R/MPRAnalyze, we performed principal components analysis to estimate broad influence of promoter (captured primarily by PC1) and position (captured by PC2) on MPRA activity. Each point represents an individual replicate, points are color-coded based on plasmid configuration, and point shape indicates position of the cloned fragment. (B) To jointly estimate effects of promoter and position across fragments in all four configurations, we included promoter and position as covariates in the MPRAnalyze model. This scatterplot displays the coefficients from Wald testing for the promoter (pink points along x-axis, *INS* promoter vs. SCP1 promoter with local FDR < 0.05) and position (orange points along y-axis, upstream versus downstream with local FDR < 0.0). PC: principal component, FC: fold-change

To systematically test all fragments in each library for enhancer activity, we used the R package MPRAnalyze (Ashuach et al. 2019), which estimates activity by comparing the RNA counts for each barcode to DNA counts for the same barcodes (**Tables S2-5**). Within each of the four different constructs, we calculated the proportion of fragments categorized as significantly active by MPRAnalyze (**Figure S2B**; FDR < 0.05): 7,724 (of 11,656 total fragments tested, 66.27%) downstream of the *INS* promoter, 5,596 (48.01%) upstream of the *INS* promoter, 6,337 (54.37%) downstream of the SCP1 promoter, and 4,680 (40.15%) upstream of the SCP1 promoter. These proportions are comparable to those calculated when using all fragments without requiring their representation across all four configurations (**Table S6**). Of the set of 11,656 fragments, only 826 (7.09%) were significantly active across all four configurations (**Figure S2C**). We also defined sets of fragments that were uniquely active in a single configuration representing a collective 2,743 fragments (23.53%): 1,114 (9.56%) downstream of the *INS* promoter, 543 (4.66%) upstream of the *INS* promoter, 686 (5.89%) downstream of the SCP1 promoter, and 400 (3.43%) upstream of the SCP1 promoter (**Figure S2C**). Taken together, these initial analyses indicate that each MPRA construct influences measured enhancer activity in a unique manner where most fragments were significantly active across only one or two constructs of the four tested.

### Quantifying promoter and position effects

We systematically quantified effects of each promoter and position on an individual fragment basis using a linear modeling approach in MPRAnalyze. We modeled the activity of each fragment with position and promoter as covariates, then performed a Wald test between each contrast (SCP1 vs. *INS* and upstream vs. downstream) to extract model coefficients (**Tables S7-8**). Overall, we identified 698/11,656 fragments (5.99%) with significant promoter effects (**Figure 2B**; *FDR* < 0.05) and 703/11,656 fragments (6.03%) with significant upstream/downstream position effects. Between the two groups, only 74 fragments display both promoter and position effects. While fragments with significant position effects were roughly divided between upstream- and downstream-bias, we observed that 73.35% of fragments with significant promoter effects were biased towards the *INS* promoter (n = 512/698; *p* = 2.2 x 10^-16^, binomial test with 55% expectation, or the proportion of fragments active with the *INS* promoter). The sequence content of the SCP1 promoter is markedly different from the *INS* promoter used here. SCP1 is an 81-bp sequence composed of four highly conserved core promoter motifs (Juven-Gershon et al. 2006): a TATA box, the initiator element (Inr), a motif ten element (MTE), and the downstream promoter element (DPE). SCP1 is known to act as a strong promoter for enhancers in conventional reporter constructs, and was used in one of the first genome-wide pooled reporter assays, STARR-seq (Arnold et al. 2013). By contrast, we used a 408-bp fragment of the *INS* promoter spanning -364 to +44 bp of the human *INS* gene (German et al. 1995) which contains a battery of pancreas-expressed TF binding sites (*e.g.,* RFX6, NEUROD1, INSM1) (Chandra et al. 2014) and designated glucose-responsive elements (Melloul et al. 2002). We hypothesized that specific sequence features of the *INS* promoter may enable regulatory compatibility with a broader subset of diabetes and related trait-associated fragments.

### Sequence features associated with position- or promoter-bias

Next, we were interested in defining specific sequence features associated with promoter- or position-driven activity bias using a set of genomic annotations, including transcription factor motifs, accessible chromatin peaks measured in human pancreatic islets, and ChromHMM-based chromatin state inferences derived from genomic profiling in islets (**Figure 3A**). Given that this set of annotations is large (∼3,000 features), we anticipate that only a small subset is associated with activity bias. To address this challenge, we used LASSO regression, an extension of linear regression that penalizes and thereby removes uninformative predictors, resulting in a simple, interpretable model. For each unique fragment sequence, we calculated overlap scores across all annotations which we standardized for use as model predictors using rank-based inverse normalization. We performed LASSO regression with 10-fold cross validation, where the annotation overlap scores were the predictors and the position or promoter bias scores (i.e., signed Wald statistics from the previous joint analysis) were the outcome variables. To account for redundancy in TF motifs, we first performed LASSO using a clustered set of 540 motifs (D’Oliveira Albanus et al. 2021) and compared to results using the full set of annotations (**Tables S9-12**). Only four features showed significant association with promoter bias, with all four enriched in fragments with *INS* promoter-biased activity versus SCP1 (**Figure 3B**). Notably, two of these were motifs for HNF family transcription factors, which play important roles in pancreatic beta cell development, differentiation, and homeostasis, and when disrupted cause maturity onset diabetes of the young (MODY), an early onset monogenic form of diabetes (Nkonge et al. 2020; Zhang et al. 2021). Concordantly, many noncoding variants associated with complex, later-onset diabetes (i.e., T2D) are enriched in HNF1 motifs (Quang et al. 2015). Additionally, the *INS* promoter itself overlaps a putative HNF1 motif; however, there is conflicting evidence as to whether HNF1A/HNF1B directly mediates *INS* transcription (Okita et al. 1999; Párrizas et al. 2001; Emens et al. 1992). Together this regulatory convergence suggests that these two TFs serve as critical regulators of transcriptional activity central to diabetes pathogenesis.

**Figure 3.**
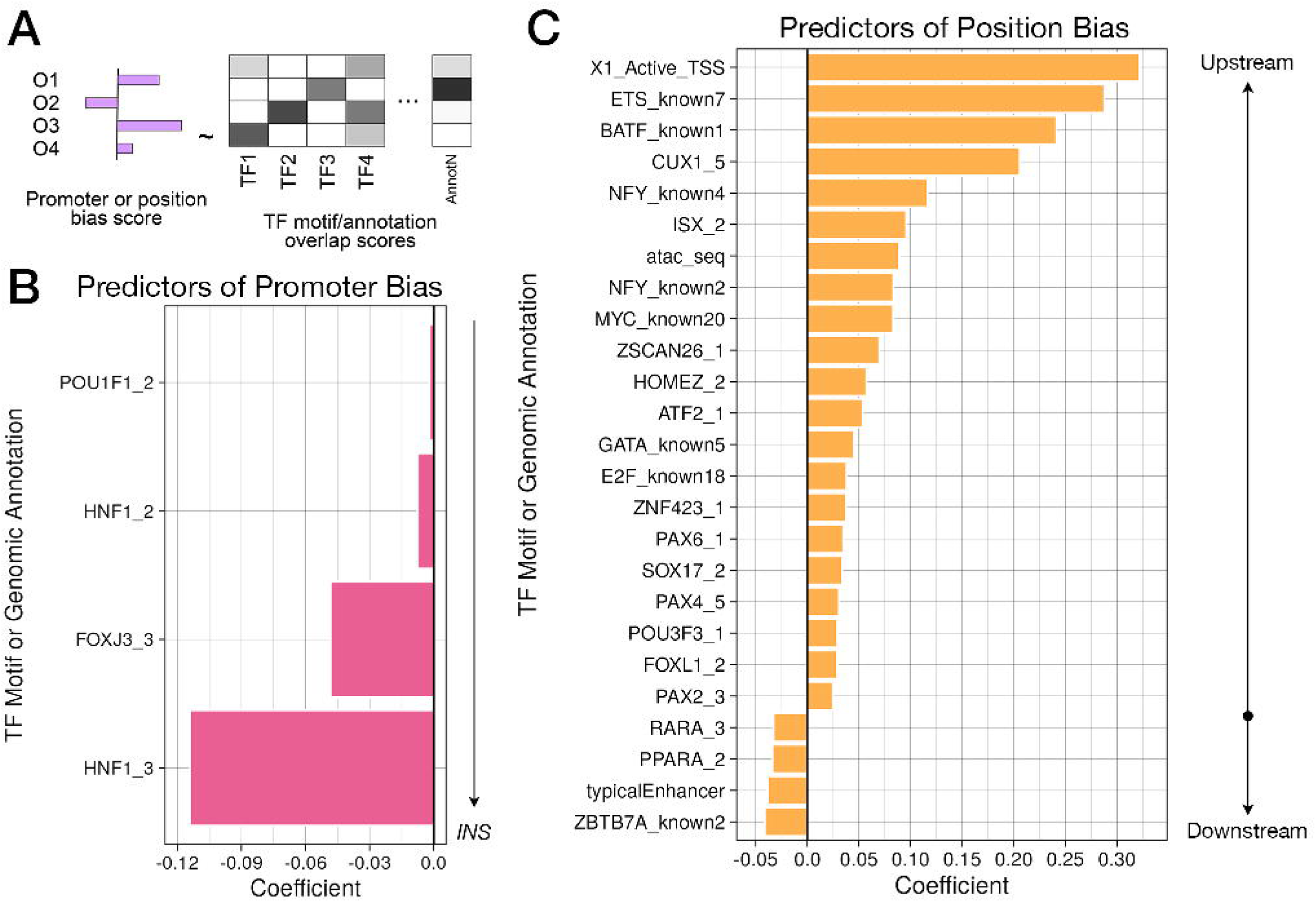
Position and promoter bias in enhancer activity is partially attributed to presence of relevant annotations. **(A)** To select sequence features associated with position or promoter bias, we performed LASSO regression with 10-fold cross validation on position or promoter bias scores (signed Wald statistics) as a function of several thousand transcription factor motifs and genomic annotations. **(B)** Significant predictors of promoter bias are displayed. With the clustered set of annotations, we identified only features enriched in fragments that are more active with the *INS* promoter. **(C)** Top 25 predictors (i.e., nonzero coefficients) associated with position bias effects are shown. Positive coefficients indicate annotations that are enriched in fragments that are more active in the upstream position while negative coefficients are enriched in fragments that are more active in the downstream position.

When performing LASSO with the unclustered set of motifs, we additionally found that motifs for NF-κB constituents (RELA/p65, NFKB) and the ubiquitously expressed activator TFs SP1 and SRF were associated with SCP1 promoter-biased fragment enhancer activity (**Figure S3A**). SP1 functions by interacting with the transcription factor II D complex (TF_II_D) (Pugh and Tjian 1990), a constituent of the RNA polymerase II preinitiation complex that binds to core sequences found in the SCP1 promoter (Louder et al. 2016). HINFP (also known as MIZF), a transcriptional repressor (Sekimata and Homma 2004; Sekimata et al. 2001), was the strongest coefficient associated with SCP1 promoter-biased activity. In addition to the annotations associated with *INS* promoter-biased activity in the reduced set, we found that several GATA family TFs and the SOX family member SRY motifs were strongly predictive of fragments’ bias towards the *INS* promoter, as was the motif for NKX6.1, another key pancreatic beta cell identity TF. Similar to *HNF1A*/*HNF1B*, coding variants in *NKX6-1* have more recently been explored as very rare causes of MODY (<1%) (Mohan et al. 2018; Taylor et al. 2013) and T2D-associated variants are also enriched in NKX6.1 motifs (Chiou et al. 2021; Khetan et al. 2021, 2018).

We applied the same approach to define features associated with upstream/downstream position bias (**Figure 3C; Figure S3B**). Of the significantly associated features, most (32/46) exhibited bias towards upstream-cloned fragments. Features predictive of upstream bias included overlap with open chromatin peaks in pancreatic islets (“atac_seq”), motifs for the transcriptional activator NF-Y (which binds to CCAAT motifs in promoters) (Dolfini et al. 2012) and several TFs (e.g., CUX1, ISX, POU family members, and PAX family members) known to bind to promoters of target genes important for developmental processes and fate specification (Vadnais et al. 2013). The strongest predictor of upstream-biased activity was presence of the “Active_TSS,” an annotation denoting active promoters as defined by ChromHMM (Ernst and Kellis 2012). Conversely, many fewer annotations were associated with downstream-biased enhancer activity, though among these included the “typicalEnhancer” ChromHMM state, which overlaps conventional enhancer marks such as p300 or H3K27ac (Vahedi et al. 2015).

### Evaluating effects of HNF1 motif perturbations on regulatory activity

To explore the putative interaction between the HNF1 motif within enhancer regions and the *INS* promoter, we designed and constructed a targeted MPRA library (**Figure 4A; Figure S1B**). We selected corresponding variants in 47 fragments with significant *INS* promoter-biased activity (FDR < 0.05) containing HNF1 motifs. To generate motif deletion or shuffled fragments, we first identified the most likely motif position using SNP-aware position weight matrix scanning. For motif deletion fragments, we removed the matched sequence and added (*n* motif bp)/2 of surrounding sequence to each end of the fragment to keep the fragment length constant. For motif shuffled fragments, we performed a dinucleotide shuffle on the matched motif sequence to disrupt the motif while maintaining the same sequence content. In cases where the variant was adjacent to the HNF1 motif (rather than contained within), we synthesized motif deletion and motif shuffled fragments with both alleles. We cloned this set of fragments into an MPRA vector upstream of the *INS* promoter then delivered this library to 832/13 cells (n = 6), as described previously.

**Figure 4.**
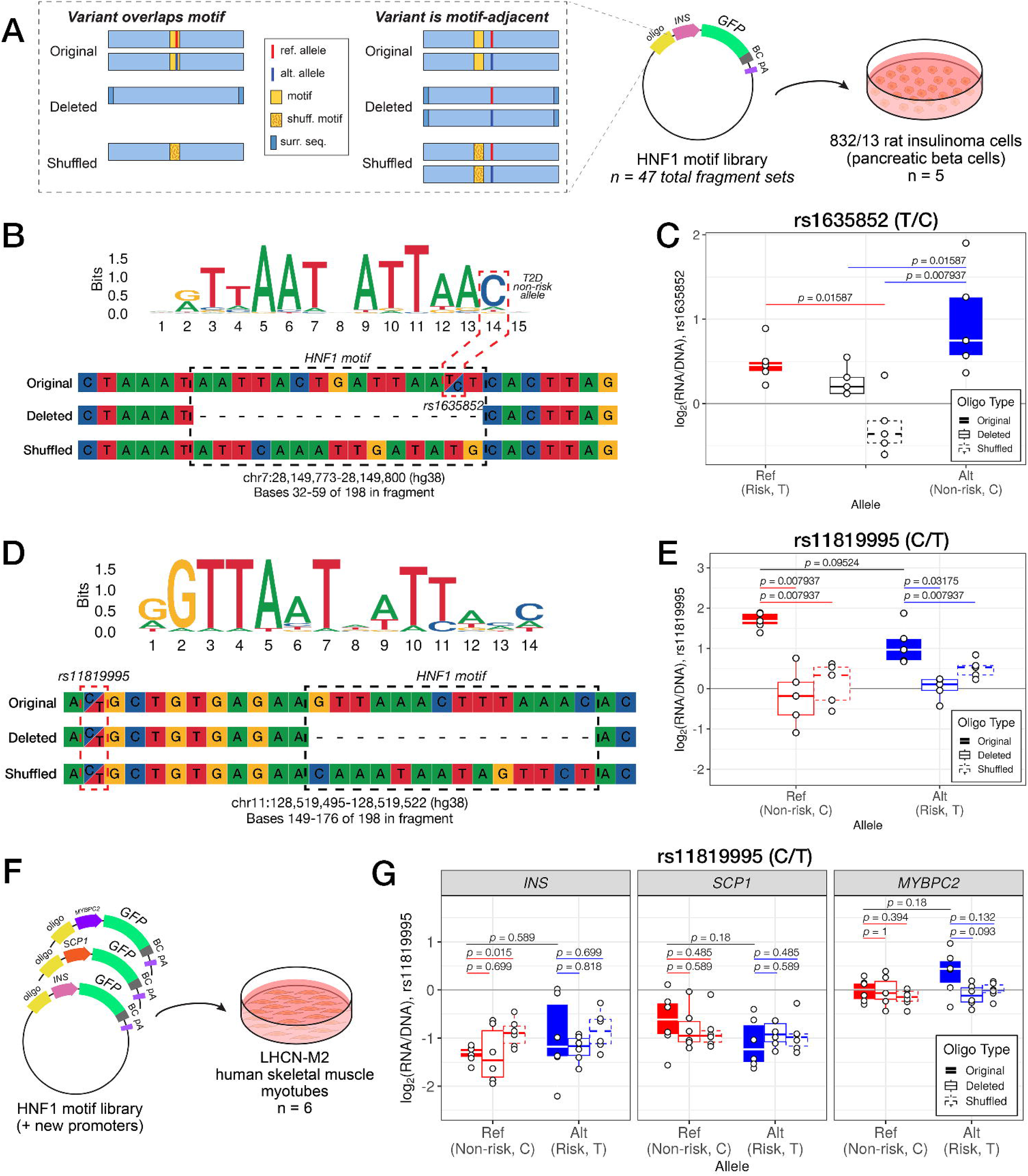
HNF1 transcription factor motifs contribute to enhancer activity near selected T2D- associated variants. **(A)** For 47 HNF1 motif-overlapping fragments with significant *INS* promoter bias effects on activity, we designed three versions: original (motif intact), deleted (motif removed and sequence adjusted), and shuffled (dinucleotide-shuffled motif). When the tested variant was adjacent to the motif, we synthesized both reference and alternate alleles for each version. For variants directly overlapping the motif, we generated only one deletion and one shuffled fragment. **(B)** We synthesized four fragments corresponding to the variant rs1635852, which overlaps an HNF1 motif at a high information content position. The T2D risk allele (T) disrupts this motif, while the non-risk allele (C) matches the consensus. **(C)** Shuffling the motif significantly decreased enhancer activity compared to intact fragments with either the risk T (Wilcoxon rank-sum test *p* = 0.016) or non-risk C allele (*p* = 0.008). Motif deletion also significantly decreased enhancer activity compared to the non-risk C allele (*p* = 0.016). **(D)** For the variant rs11819995, located 11 bp upstream of an HNF1 motif, we synthesized six fragments. **(E)** Deletion of the motif significantly decreased enhancer activity for both the reference (C, non-risk) and alternate (T, risk) alleles (*p* for reference allele = 0.008 and for alternate allele = 0.03). Shuffling the motif likewise reduced activity for both alleles (*p* = 0.008). **(F)** To assess context-specific effects, we cloned these fragments into MPRA vectors with the SCP1 or skeletal muscle-specific *MYBPC2* promoter and delivered all three to LHCN-M2 human skeletal muscle myotubes (n = 6). **(G)** When paired with the *INS* promoter, the shuffled rs11819995-containing fragment showed increased activity relative to the original fragment (*p* = 0.015); however, none of the fragments containing rs11819995 functioned as enhancers in LHCN-M2 myotubes, regardless of promoter context. Overall, their activity is highest when paired with the skeletal muscle-specific promoter.

Of the 47 fragment sets we synthesized and tested, 31 sets had at least one fragment with significant activity (FDR < 0.05) and were retained for downstream analysis (**Table S13**). One such variant was rs1635852, which is found in the first intron of *JAZF1*, a well-characterized transcriptional regulator of the pancreatic beta cell response to metabolic stress (Kobiita et al. 2020). This variant falls in a high-information content position of the HNF1 consensus motif (JASPAR record MA0046.2), where changing from the reference allele (T, risk allele for T2D) to the alternate allele (C, non-risk) is predicted to strengthen the motif (**Figure 4B**). We did not observe a statistically significant allelic difference in activity between the reference and alternate alleles, likely owing to high variance in the alternate allele fragment (Wilcoxon rank-sum test *p* = 0.22); however, the effect is in the expected direction (Fogarty et al. 2013). We did observe that shuffling the HNF1 motif led to a significant decrease in activity compared to either the risk (Wilcoxon rank-sum test *p* = 0.01587) or non-risk allele fragments (**Figure 4C**; Wilcoxon rank-sum test *p* = 0.007937). Deleting the HNF1 motif led to significantly decreased activity compared to the non-risk allele alone (Wilcoxon rank-sum test *p* = 0.01587). To evaluate the extent to which these effects were restricted to the *INS* promoter, we also tested these fragments with the SCP1 promoter. We observed no significant differences in activity based on HNF1 motif status (**Figure S4A**). A previous study reported higher activity for the alternative compared with the reference allele using reporter and DNA affinity assays (Fogarty et al. 2013), consistent with the direction of effect we observed. Interestingly, this study concluded that regulatory activity at this locus was mediated by differential interaction with another key transcription factor expressed in beta cells, PDX1. In contrast to the HNF1 motif, the rs1635852 risk T allele is slightly preferred over the non-risk C allele in the PDX1 motif (JASPAR record MA0132.2). Because our scrambled HNF1 motif oligo also partially disrupts the PDX1 motif, this assay may also be detecting changes to PDX1 binding. Our results point to a possible complementary regulatory mechanism at this locus mediated by HNF1A/B. We again observed HNF1 motif-dependent activity at rs11819995, a T2D signal variant recently identified in a multi-ancestry GWAS by the DIAMANTE consortium (Mahajan et al. 2022). This variant is located in the first intron of the canonical *ETS1* transcript and overlaps an alternative transcription start site active in pancreatic islets (Varshney et al. 2017; Mahajan et al. 2022). The 14-bp HNF1 motif matched by position weight matrix scanning (*GTTAAACTTTAAAC*) starts 11 bp downstream of the variant in the fragment sequence (**Figure 4D**). Here, we observed a modest allelic effect in fragments with intact HNF1 motifs (Wilcoxon rank-sum *p* = 0.09524). Deleting the HNF1 motif significantly decreased the regulatory activity of fragments, in the presence of either the reference (C, non-risk allele for T2D; Wilcoxon rank-sum test *p* = 0.007937) or alternate allele (T, risk; Wilcoxon rank-sum test *p* = 0.03175) of this variant (**Figure 4E**). Additionally, shuffling the HNF1 motif in fragments with either allele decreased activity in comparison to the original fragment (Wilcoxon rank-sum test *p* = 0.007937). As with the previously described variant, we tested fragments spanning rs11819995 in a construct with the SCP1 promoter. None of these fragments were differentially active based on whether the motif was intact, deleted, or shuffled (**Figure S4B**).

To further explore HNF1 motif-*INS* promoter interaction, we considered two related but distinct questions: (1) does perturbing the HNF1 motif have similar effects when cloned in proximity to a different tissue-specific promoter, and (2) is the interaction between the HNF1 motif and *INS* promoter restricted to a pancreatic beta cell line? To these ends, we cloned the same library of original, deleted, or shuffled fragments in one of two new promoter contexts, with either the SCP1 promoter or with the promoter for the skeletal muscle-specific gene *MYBPC2*. We then delivered these two libraries along with the *INS* promoter MPRA library to the LHCN-M2 human skeletal muscle myoblast line (n = 6) after differentiation to myotubes (**Figure 4F**). In this line, *MYBPC2* is abundantly expressed, and *INS* is transcriptionally silent.

In the LHCN-M2 cell line, none of the HNF1 motif fragment sets displayed significant enhancer activity (log_2_(RNA/DNA) > 0 and FDR < 0.05) when paired with any of the three different promoters (**Figure 4G**). Additionally, we observed that their activity was modulated by promoter context, where the sequences tended to be more active when cloned with the muscle-active *MYBPC2* promoter versus SCP1 or *INS1*. Finally, we observed only one instance in which disrupting the HNF1 motif led to modulated regulatory activity. Compared to the original fragment, scrambling the HNF1 motif caused a modest increase in enhancer activity when paired with the *INS* promoter (Wilcoxon rank-sum test *p* = 0.015); however, HNF1A and HNF1B are not expressed in skeletal muscle, so enhancer activity of this fragment is likely driven by a separate transcription factor. In all other cases across all three promoter pairings, we no longer observed any dependence upon the HNF1 motif. Taken together, this indicates HNF1A/B-orchestrated regulatory activity is beta cell-specific.

## Discussion

Massively parallel reporter assays (MPRAs) enable large-scale variant-to-function studies of cis-regulatory elements. However, MPRA implementations differ in key technical aspects including the cloning and configuration of the reporter construct, whether it is integrated or episomal, and which basal promoter it carries. Differences in sequence and cellular context can influence the activity of the cloned fragments (Klein et al. 2020; Bergman et al. 2022); however, given these screens’ scale, it is not routine practice to examine how each of these factors influence individual fragments’ activity or how they may modulate the effects of genetic variation.

In this study, we used a modular MPRA construct to explore the effects of sequence context and proximity upon regulatory elements’ activity. We found that for a subset of disease-associated genomic sequences, enhancer activity was sensitive to proximity and identity of the promoter used, independent of overall activity. We found specific genomic annotations that were associated with these biases and noted that direction of effect was concordant with those interpretations. For instance, fragments annotated as proximal transcription start sites by ChromHMM tended to be more active when cloned upstream, and conversely, those annotated as enhancers were more active when cloned downstream. Of note, we discovered that sequences preferentially active with the human insulin (*INS*) promoter were enriched in motifs for the important pancreatic beta cell transcription factors including HNF1A/B and NKX6.1. In a follow-up MPRA, we demonstrated the necessity of HNF1A/B motifs for activity of a subset of these fragments and confirmed that the dependence upon those motifs was specific to both the cell type and the paired promoters.

Our observation that some sequences’ activity depends upon proximity to a promoter is consistent with many previous reports. Enhancers are known to work at a range of distances, including 1,000s of kb away from their targets. Here, we identified that sequences that were more active when placed downstream (distal to) the promoter and reporter gene tended to be enriched for features of typical enhancers while those that were more active in the upstream position (proximal) contained promoter-like elements including enrichment for active TSSs. It is important to note that the MPRA construct we used in this study to explore positional effects is distinct from the STARR-seq vector in that fragments are cloned downstream of the poly-A signal and thus are not transcribed (Klein et al. 2020; Das et al. 2023). Therefore, we do not suspect that interaction with RNA-binding proteins or accelerated RNA decay influenced the results observed here.

Previous studies using reporter assays have indicated that core promoter sequences display some level of tissue-specificity (Mattioli et al. 2019), where observed activity in an MPRA vector is strongly correlated with endogenous transcriptional activity from the same loci. This has also been observed by others working to design minimal synthetic promoters that can function in a context-dependent manner, including with respect to cell type (Zahm et al. 2024). Together with our work, this suggests that promoter sequences contain a unique syntax that interacts with a cell’s regulatory milieu to produce highly tuned gene expression beyond the standard elements common to many promoters (e.g., TATA box, downstream promoter element, etc.). It is important to note that we used a well-delimited promoter for the highly cell type-specific human insulin gene that was previously defined through deletion mapping and motif analysis (German et al. 1995; Melloul et al. 2002). Selecting suitable promoters for other cell types will require a similar level of inspection.

Though not explicitly considered in the work presented here, the choice of using an episomal or integrated MPRA vector has also been examined previously (Inoue et al. 2017; Klein et al. 2020; Guzman et al. 2023). Here, we used an integrated construct to compare activity at a specific locus in the LHCN-M2 human skeletal muscle cell line compared to an electroporated episomal construct in the 832/13 rat beta cell line (**Figure 4D-E**), due to technical limitations that make electroporation or transfection of differentiated LHCN-M2 cells unfeasible. While we see differences in enhancer activity across the two lines, we believe that this is a biological distinction and not a methodological one. An additional source of concern for episomal MPRAs is preferential usage of the bacterial origin of replication (ORI) compared to the SCP1 promoter (Muerdter et al. 2018; Klein et al. 2020). Consequently, researchers have advised to use other synthetic promoters (e.g., minP) or ORI-less versions of the MPRA backbone to avoid this issue. Given the systematic discrepancies measured between the two methods in the studies referenced above, this design choice may also require careful forethought.

Although our initial, large MPRA library contained both reference and alternate alleles across a large set of T2D-associated variants, it was not cloned at sufficient complexity to allow for sensitive detection of allelic effects, which are often subtle. We therefore focused our analyses on fragment-level activity across position and promoter contexts. This approach complements prior allele-focused MPRAs, such as one published in 2021 by Khetan and colleagues (Khetan et al. 2021) that characterized over 2,000 T2D-associated variants. Given that the number of T2D-associated genomic loci has more than quadrupled since these libraries were constructed (Suzuki et al. 2024; Mahajan et al. 2022; Vujkovic et al. 2020), it is a high priority to develop new MPRA libraries to profile the expanded set of loci.

The work presented here indicates that in MPRA experiments, proximity to the promoter and the choice of promoter each have measurable impacts which may vary across different fragments. Though it takes consideration to select appropriate promoters for a cell type of interest, we show that for pancreatic islet beta cells using the *INS* promoter was sufficient to capture regulatory activity at many T2D-associated loci. Given the inherent sparsity of MPRA datasets, employing these libraries across a variety of design configurations and cell types will enhance our ability to understand regulatory processes that are disrupted in complex diseases.

## Methods

### Cell culture

We obtained INS-1 832/13 rat insulinoma cells from Dr. Christopher Newgard (Sarah W. Stedman Nutrition and Metabolism Center, Duke University, Durham, NC) and LHCN-M2 human skeletal muscle myoblasts from Evercyte. We cultured 832/13 cells in RPMI-1640 containing 10% fetal bovine serum, 0.05 mM β-mercaptoethanol, 1 mM sodium pyruvate, 2 mM L-glutamine, 10 mM HEPES, and 1,000 U/mL penicillin/streptomycin. We maintained LHCN-M2 cells on 0.1% gelatin-coated flasks in 4:1 DMEM (4.5 g/L glucose, glutamine, bicarbonate):Medium 199 (with bicarbonate) containing 15% FBS, 20 mM HEPES, 30 ng/mL zinc sulfate, 1.4 ug/mL vitamin B12, 55 ng/mL dexamethasone, 2.5 ng/mL recombinant human hepatocyte growth factor, 10 ng/mL basic fibroblast growth factor, and 600 U/mL penicillin/streptomycin.

### Large (first) MPRA library design, construction, and delivery

We compiled regions of interest for the large MPRA library from loci associated with type 2 diabetes (T2D) and related metabolic traits, drawing on signal variants reported by the DIAMANTE Consortium (Mahajan et al. 2022) and signal variants for HbA1C, fasting insulin adjusted for BMI, fasting glucose adjusted for BMI, and 2-hr glucose adjusted for BMI reported by the MAGIC Consortium (Chen et al. 2021). To provide broader disease context, we also included loci for non-T2D traits including rheumatoid arthritis, systemic lupus erythematosus, and type 1 diabetes. We used UK Biobank imputed genotypes to calculate linkage disequilibrium and fetch proxies with an R^2^ greater than 0.8. We removed any variants that were not SNVs (i.e., indels, CNVs) resulting in a set of 13,226 sites. For each site, we designed 198-bp fragment sequences in three positional configurations relative to the focal base: offset left (position 50 of 198 bp), centered (position 99), or offset right (position 149). In cases where a variant had two alternate alleles, we designed both sets of sequences. We extracted all sequences from the hg19 reference genome. We added 16-bp flanking adapter sequences for PCR amplification and cloning. We obtained a pool of 79,511 full-length 230-bp fragments from Agilent.

We constructed the large MPRA library as described previously (Varshney et al. 2021) (**Figure S1A**). We added homology arms via PCR, then cloned amplified fragments into either a KpnI- or EcoRV-digested (upstream or downstream, respectively) MPRA backbone via HiFi assembly. We transformed this first assembly into electrocompetent 10-beta cells. We post-barcoded by digesting the plasmid library with PmeI then used HiFi assembly to insert 16-bp random barcodes at the PmeI restriction site. We transformed this reaction into electrocompetent 10-beta cells and prepared final plasmid libraries for transfection using the ZymoPURE Plasmid Maxiprep Kit.

For the first library, we electroporated 50 ug of plasmid into 25 million 832/13 rat insulinoma cells for each biological replicate and lysed cells with TRIzol 24 hours later. After phase separation, we used the Direct-zol RNA Miniprep Kit to isolate total RNA. We enriched for mature mRNA transcripts with fragment(dT) beads then treated 2 ug of mRNA with DNase I to remove plasmid or genomic DNA contamination. We reverse transcribed 1 ug of mRNA using SuperScript III with a custom primer containing a 6-bp unique molecular identifier (UMI) targeting transcripts derived from the MPRA plasmids (**Table S14; Figure S1C**). We treated cDNA samples with DpnI to eliminate any residual plasmid DNA.

### Motif (second) MPRA library design, construction, and delivery

We constructed the motif MPRA library using a modified version of the methods detailed in Tewhey, et al. 2016. While the cloning strategy differed slightly from what we used for the first library, the assay design and functional readout between the two are comparable (**Figure S1B**). We selected fragment sequences from the first library with strong *INS* promoter-biased activity that contained HNF1 motifs. We obtained fragments spanning the same amount of genomic sequence as the previous library but containing different adapters (IDT). We added barcodes and a promoter cloning scaffold via PCR, then cloned the amplified fragments into a modified pMPRA1 vector (a gift from Tarjei Mikkelsen; Addgene #49349) using Golden Gate assembly with PaqCI. We digested this assembly with SfiI to remove empty backbones then transformed into electrocompetent 10-beta cells. We inserted clonal promoter fragments using Golden Gate assembly and digested with AsiSI to remove promoterless constructs. We transformed this final assembly into electrocompetent 10-beta cells, expanded in 150 mL cultures, and isolated plasmid using the ZymoPURE Plasmid Maxiprep Kit.

For the second library, we electroporated 30 ug of plasmid into 30 million 832/13 cells. We lysed cells 24 hours later with Buffer RLT Plus and isolated total RNA with the Qiagen RNeasy Midi Kit. We reverse transcribed 30 ug of DNAse I-treated total RNA using SuperScript IV (**Figure S1B**). We performed a cDNA amplification PCR and sample indexing with all cDNA made.

For use with the LHCN-M2 skeletal muscle myocyte cell line, we ported the assembled MPRA block (fragment, barcode, promoter, GFP) to a lentiviral transfer vector (a gift from Nadav Ahituv; Addgene #137725) via restriction cloning (Gordon et al. 2020; Inoue et al. 2017). The University of Michigan Viral Vector Core produced infectious lentiviral particles with this transfer vector and third generation lentiviral plasmids in HEK293T cells. Per replicate, we infected 4 x 10^6^ LHCN-M2 human skeletal myoblasts with our MPRA library at an MOI of ∼10. After infection, we passaged the cells for one week to remove any unincorporated virus or contaminating transfer plasmid, then differentiated the cells for one week. We isolated RNA and gDNA from each replicate using the Qiagen AllPrep DNA/RNA mini kit.

For all samples generated through use of the second set of MPRA libraries, we reverse transcribed RNA into cDNA with a GFP-specific primer containing a 6-bp UMI, then constructed indexed sequencing libraries for both the cDNA and gDNA libraries using Illumina-compatible primers (**Table S15**).

### Sequencing data collection

We sequenced pairing and cDNA/input sequencing libraries for the initial set of MPRA studies on an Illumina Hiseq. We sequenced libraries generated for the second set of MPRA on an Illumina Novaseq 6000. For both sets of studies, we received paired-end 150-bp reads.

To create the barcode-fragment pairing dictionary, we used a custom pipeline. First, we merged the paired-end 150-bp reads using bwa v0.7.17 then extracted barcode and fragment sequences using cutadapt v4.3 (Li and Durbin 2009; Martin 2011). To align the merged fragment reads against our reference FASTA file, we used minimap2 v2.24 (Li 2018). After filtering for mapping quality and sequencing depth, we created a final table with fragment-barcode pairs and removed any duplicate barcodes.

We processed MPRA sequencing data using a custom pipeline that quantifies barcode counts while accounting for possible sequencing errors. We identified reads with the barcode sequence and trimmed constant sequences using cutadapt v4.3. We used UMI-tools v1.1.2 to identify and cluster UMIs, then used starcode-umi to cluster UMI-barcode and perform deduplication (Zorita et al. 2015; Smith et al. 2017). Finally, we used starcode to cluster and count barcodes.

### Activity estimation and joint modeling

Prior to model fitting, we merged cDNA and input barcode counts with the corresponding pairing dictionary requiring an exact match. For both MPRA libraries, we used the R package MPRAnalyze to estimate enhancer activity and promoter or position bias (Ashuach et al. 2019). This modeling approach fits two separate models for cDNA and input counts, then relates the two to yield a per-fragment transcription rate. For all analyses, resulting enhancer activity scores were converted from a natural log scale to binary log scale, though values are reported on both scales in Tables S2-5.

For the first MPRA library, we used this method to estimate enhancer activity within each construct type and then performed *post hoc* comparisons outside of the analysis framework. To evaluate quantitative effects of promoter and position on fragment activity in the first MPRA library, we jointly estimated enhancer activity across all four constructs and included promoter and position as covariates.

For the second MPRA library, we used this method to estimate enhancer activity across the entire library within the two cell types (832/13 and LHCN-M2) and nominate candidate fragments for further inspection. Then, on a per fragment basis, we compared activity differences across fragment types using a Wilcoxon rank-sum test.

### LASSO regression to select features associated with promoter or position bias

Here we used least absolute shrinkage and selection operator (LASSO) regression to model MPRA activity bias (promoter or position) and select genomic features associated with this bias (Tibshirani 1996). LASSO regression uses a penalty term to shrink coefficients of less informative variables to zero, thereby removing them from the model and enhancing interpretability. We quantified activity bias by using the Wald statistics from our joint estimates of enhancer activity and multiplying them by the sign of the log-fold change to indicate direction of bias (INS versus SCP1 for promoter and up-versus downstream for position). As features in our models, we used a set of annotations and TF motifs described previously (Varshney et al. 2021; D’Oliveira Albanus et al. 2021). For each fragment, we quantified motif occurrences or annotation overlap. To generate overlap scores that could be used as parameters for LASSO regression and compared uniformly across annotations, we inverse-normalized the -log_10_(p-value) for each motif or annotation overlap using R/RNOmni (McCaw et al. 2020). We ran the regression using R/glmnet with default parameters (α = 1 corresponding to LASSO) and 10-fold cross validation using the signed Wald statistic as the outcome and annotation scores as the predictors (Friedman et al. 2010). We performed regression analyses for the promoter and position bias scores separately and glmnet automatically determined the appropriate lambda value by minimizing the mean cross-validated error.

## Data and code access

All processed sequencing data generated in this study have been submitted to the NCBI Gene Expression Omnibus (GEO) under accession numbers GSE279057 (first library, GSE279071 (second library), and GSE247455 (LHCN-M2 data). Corresponding raw sequencing data is available on the NCBI Sequencing Read Archive under accession numbers PRJNA1142481 and PRJNA1142558. Complete results files are available as part of this manuscript as Supplementary Tables.

Custom scripts used to preprocess and analyze data from the second MPRA library and other auxiliary analyses are available at https://github.com/adelaidetovar/modular-t2d-mpra.

## Acknowledgements

The authors would like to acknowledge members of the Kitzman and Parker labs (University of Michigan) for their critical feedback. The authors would also like to thank the University of Michigan Viral Vector Core for producing lentivirus and the University of Michigan Advanced Genomics Core for sequencing services.

## Contributions

Conceptualization by A.T., Y.K., J.O.K., and S.C.J.P. Fragment design and MPRA library cloning by Y.K., A.T., and K.N. Data processing and analysis by A.T., Y.K., and M.B., with contributions from A.V. A.T. wrote the manuscript with input from and editing by J.O.K. and S.C.J.P.

