## Supplemental Figures for "Using a modular massively parallel reporter assay to discover context-dependent regulatory activity in type 2 diabetes-linked noncoding regions"

**Supplementary Figures and Figure Legend**

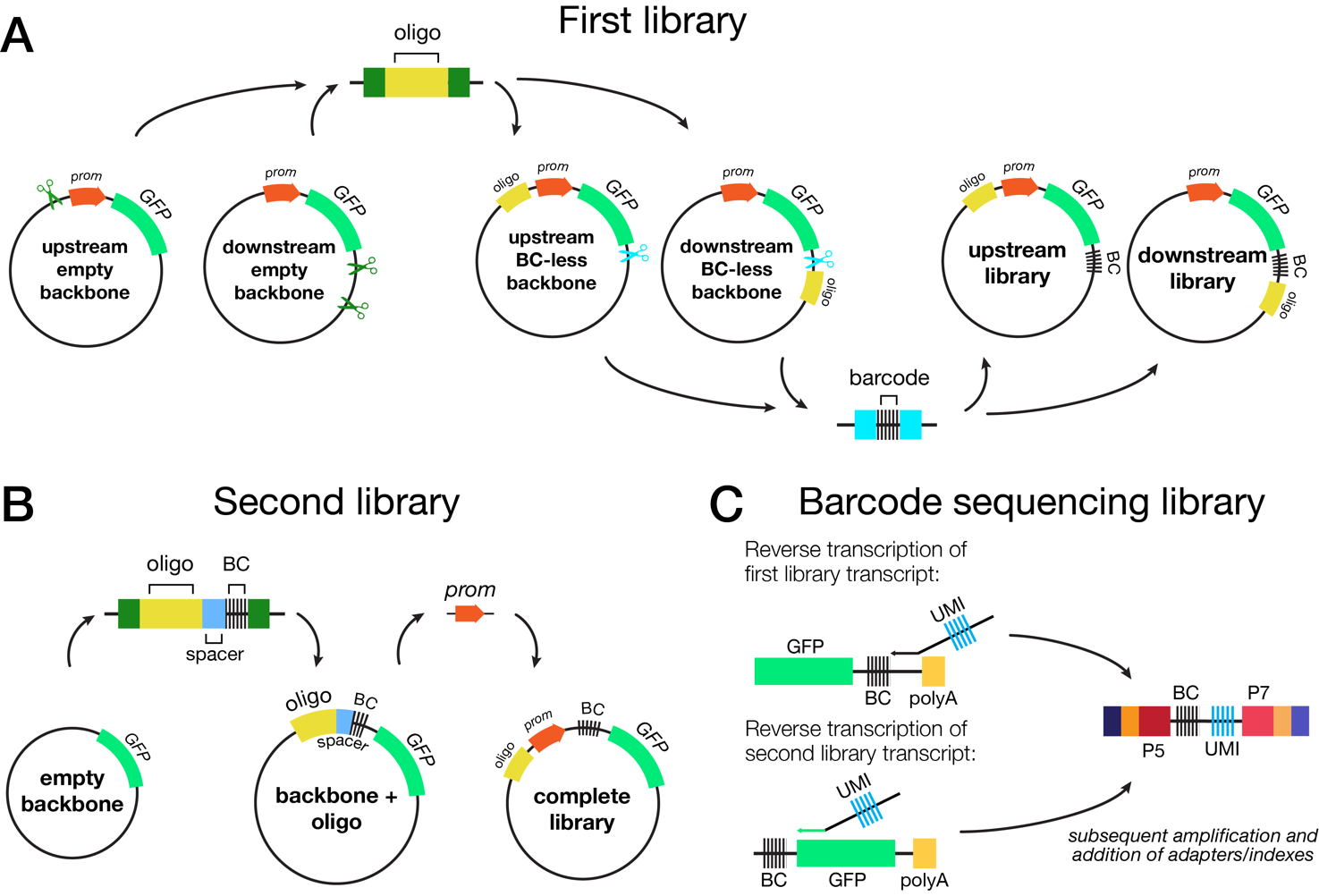

**Figure S1. Construction of MPRA plasmid libraries and barcode sequencing libraries.** (A) Construction process for the first MPRA library. We inserted oligo sequences via Gibson assembly into KpnI- or EcoRV-digested (upstream or downstream, respectively) backbones that contained either the SCP1 or human insulin (*INS*) promoter. We digested the resulting barcode-less plasmids with PmeI and inserted barcodes. (B) Construction process for the second MPRA library. We PCR-barcoded oligo sequences and inserted them into the backbone via PaqCI-mediated Golden Gate cloning. We inserted the SCP1 or *INS* promoter in a spacer sequence between the oligo and barcode via BsaI-mediated Golden Gate cloning. (C) To construct barcode sequencing libraries using mRNA extracted from cells, we performed reverse transcription using a custom primer containing a unique molecular identifier (UMI) to correct for PCR duplication in sequencing data. Primers were designed separately for each of the two MPRA types to accommodate flanking sequence differences. Subsequently we amplified the cDNA and added sequencing indexes/adapters via PCR.

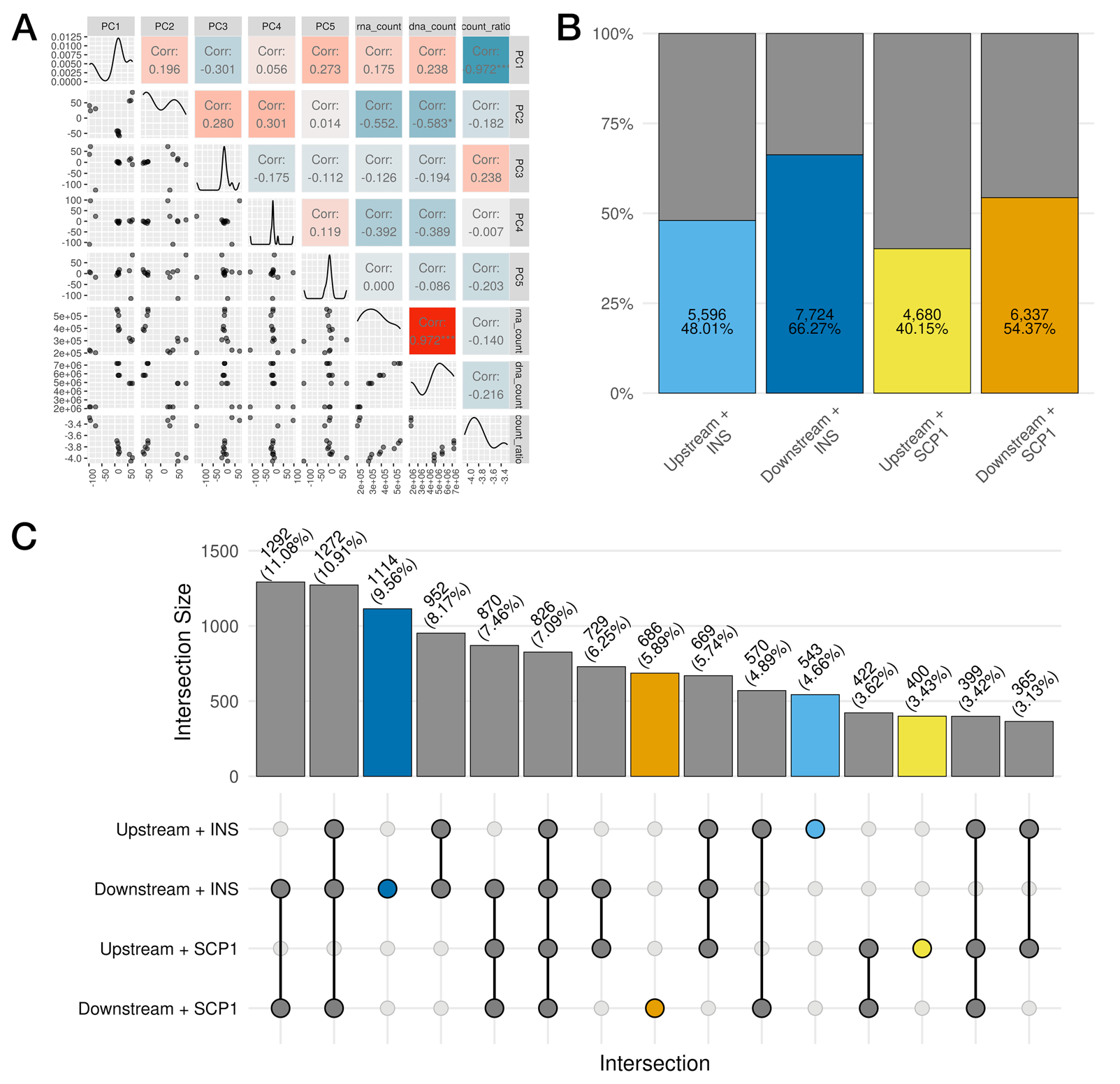

**Figure S2. MPRA plasmid configuration modestly influences overall activity of tested fragments and specific sets of significantly active (FDR < 0.05) fragments.** (A) Pairwise Spearman correlation heatmap for the first five principal components (from analysis presented in **Figure 2A**), library size-normalized RNA and DNA counts, and ratio between normalized RNA and DNA counts. (B) Stacked barplot showing the number and proportion of fragments that are significantly active (FDR < 0.05) in each plasmid configuration. Percentages are calculated using 11,656 (total number of common fragments) as the denominator. (C) UpSet plot displaying intersections (shared and unique) between the sets of fragments that are active in each plasmid configuration. Sets are depicted in the matrix on the bottom, where nodes represent individual configurations (labeled on the left) and edges connect the nodes to form intersections. The bar chart on the top shows the size of each intersection as both the number and proportion of fragments that are significantly active (FDR < 0.05).

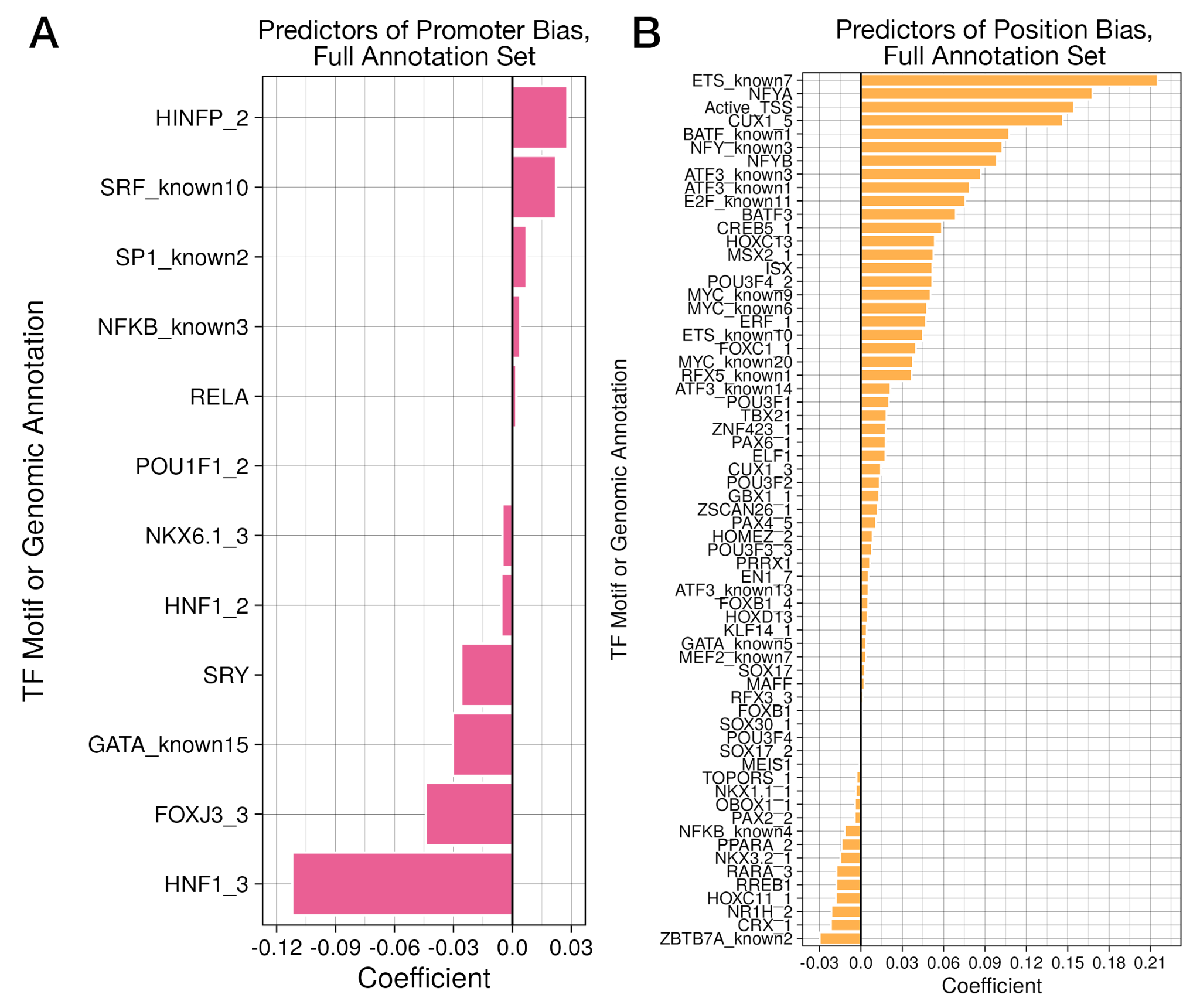

**Figure S3. LASSO regression analysis largely identifies the same set of features associated with position or promoter bias when using an expanded set of transcription factor motifs.** (A) Full set of significant predictors of promoter bias are displayed as a companion to **Figure 3B**. We identified largely the same set of coefficients. One notable addition is NKX6.1, another beta cell-specific transcription factor. (C) Full set of predictors of position bias effects are shown. The ‘activeEnhancer’ chromatin annotation is no longer listed, though ‘X1_Active_TSS’ is still among the strongest predictors of upstream positional bias.

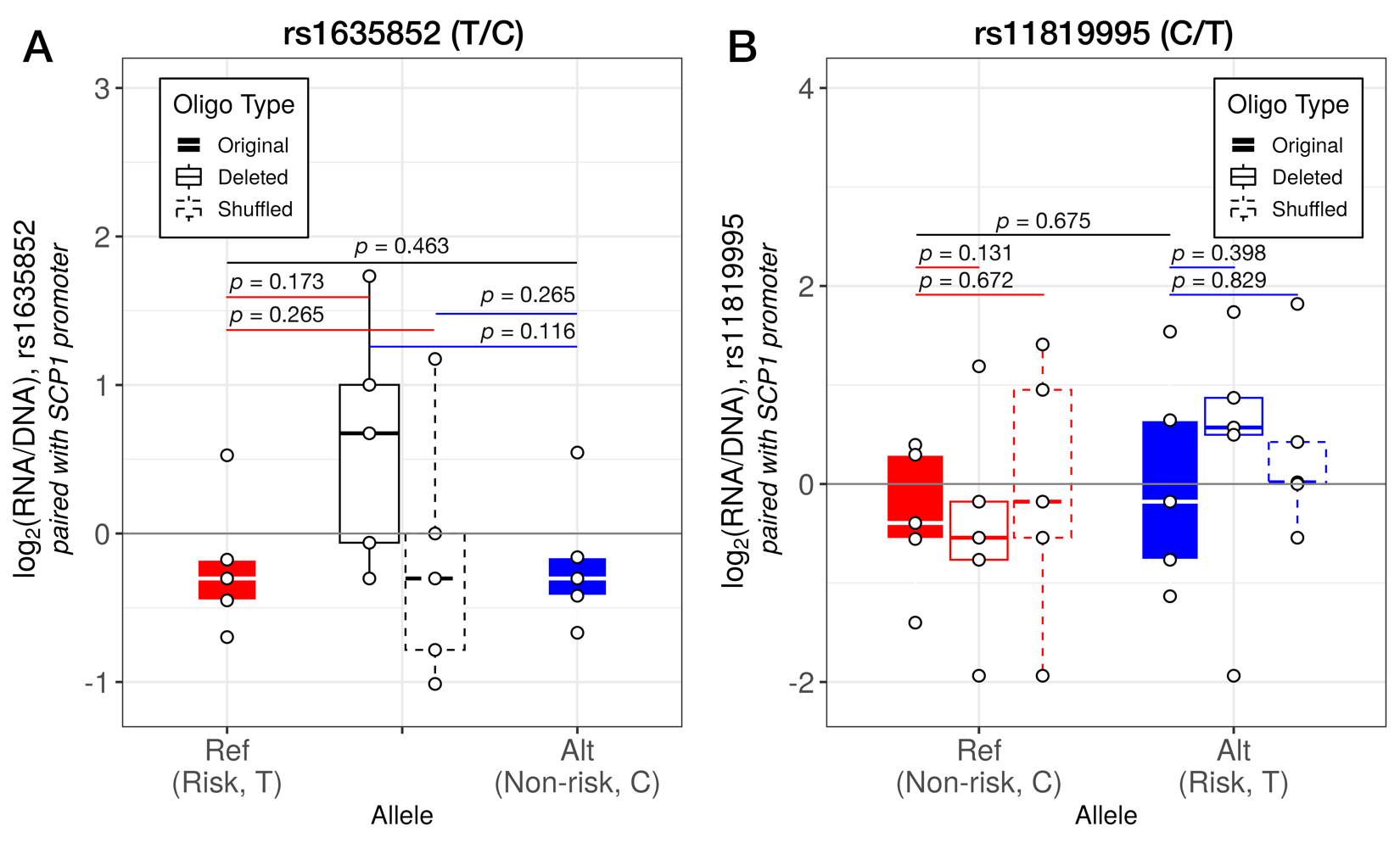

**Figure S4. Enhancer activity is minimal for variants near** **HNF1 transcription factor motifs when paired with the SCP1 promoter.** For fragments described in **Figure 4**, we also examined activity when paired with the synthetic SCP1 housekeeping promoter. **(A)** Manipulating the HNF1 motif overlapping the T2D risk variant rs1635852 has no significant impact on enhancer activity, nor is there an allelic difference in activity between the risk (T, reference) and non-risk (C, alternate) alleles. **(B)** Similar to the previous variant, none of the fragments for the variant rs11819995 have significant activity. Additionally, manipulating the HNF1 motif near this variant has no significant effect on activity compared to the original, motif-intact versions of these fragments (Wilcoxon rank-sum test *p* values reported, n = 5).
